# 40 Hz Steady-State Visually Evoked Potentials Recovered During Oscillating Transcranial Electrical Stimulation

**DOI:** 10.1101/2024.06.21.599984

**Authors:** Laura Hainke, Manuel Spitschan, Josef Priller, Paul Taylor, James Dowsett

## Abstract

**Objective:** Transcranial Electrical Stimulation and Visual Stimulation in the gamma band (30-100 Hz, especially 40 Hz) are increasingly used to study and even enhance human cognition. Combining both techniques would be of scientific and clinical value, provided that Steady-State Visually Evoked Potentials (SSVEPs) are measured concurrently to determine any neuronal effects of the electrical brain stimulation. This poses a substantial methodological challenge. We aimed to demonstrate that recording visually evoked 40 Hz activity with EEG during electrical brain stimulation is possible and to explore potential interactions.

**Approach:** We tested if electrical and visual stimulation might interact depending on which brain areas are electrically stimulated (Experiment 1; *N*=25) and how closely the respective frequencies match (Experiment 2; *N*=25). Experiment 3 (*N*=25) assessed how effectively the data processing pipeline can mitigate electrical artefacts and recover real evoked neuronal activity. SSVEPs were processed and analysed in the time domain using an optimised adaptive template subtraction approach.

**Main Results:** 40 Hz SSVEPs were successfully recorded during frequency-matched electrical stimulation applied between central and occipital regions. Waveform correlations revealed that SSVEPs from combined visual and electrical stimulation were more similar in shape to baseline SSVEPs from visual stimulation alone than to control data from electrical stimulation alone. Accordingly, during combined stimulation, the recovered signals were stronger in amplitude than the electrical control data. We found no evidence of interactions between electrical and visual stimulation.

**Significance:** We demonstrated that 40 Hz SSVEPs can be reliably measured with EEG during frequency-matched electrical brain stimulation, distinguishing neuronal activity from electrical or physiological confounds. This method enables fundamental and clinical researchers to combine rhythmic sensory and electrical stimulation in the gamma band and concurrently quantify neuronal electrophysiological effects.

## Introduction

Visual Stimulation (VS) and Transcranial Electrical Stimulation (TES) can be applied rhythmically to safely and effectively modulate neuronal oscillations. Oscillations in the gamma band (30-100 Hz, especially at 40 Hz) are of particular interest due to their role in perception and memory (Herrmann et al., 2010) and because they are disrupted in disorders such as Al zheimers Disease (Güntekin et al., 2022). By modulating gamma activity, both VS and TES applied in this frequency band are promising approaches to causally investigate related cognitive functions (Hanslmayr et al., 2019) and to intervene against cognitive decline (Guan et al., 2022; Nissim et al., 2023; Shu et al., 2024; Strüber & Herrmann, 2020; Traikapi & Konstantinou, 2021). Since audiovisual stimulation yields larger effects than visual alone (Blanco-Duque et al., 2023), combining visual and electrical stimulation could potentially also increase effects, but this approach is untested. Importantly, neuronal responses would need to be measured during stimulation, as the modulation of neuronal activity is necessary for studying gamma-related cognitive functions (Hanslmayr et al., 2019) and for clinical efficacy (Blanco-Duque et al., 2023).

Mechanistically, rhythmic VS at a given frequency evokes oscillatory brain activity measurable in the form of Steady-State Visually Evoked Potentials (SSVEPs), especially in the visual cortex (Bayram et al., 2011; Herrmann, 2001). By contrast, oscillatory TES modulates cortical excitability periodically and entrains neuronal firing to the applied electrical current (Groppa et al., 2010; Herrmann et al., 2013). Therefore, if oscillatory TES is applied to the visual cortex and at a frequency matching VS, we posited that the visually evoked activity could resonate with and be enhanced by TES. It follows that to investigate this potential interaction between VS and TES, brain activity should be measured online, during stimulation. However, this poses several challenges. First, Magneto- or Electroencephalography (M/EEG) recorded during TES is naturally contaminated by electrical artefacts, which are several orders of magnitude larger than the neuronal signal of interest (Kasten & Herrmann, 2019; Noury et al., 2016). Second, neuronal gamma power is relatively low and can be confounded with ocular or muscular artefacts (Hipp & Siegel, 2013). Third, stimulating visually and electrically at the same frequency makes it hard to distinguish whether the resulting rhythmic signal rather represents evoked brain activity or residual electrical artefacts.

Prior research has mostly focused on mitigating TES artefacts in EEG data, which remains challenging (Kasten & Herrmann, 2019). One approach is based on subtracting a template of the TES artefact from EEG segments; outcomes are quite sensitive to stimulation protocols and analysis parameters (Dowsett & Herrmann, 2016). A separate line of work reports the benefits of combined TES and VS, but without online electrophysiological measurements (Li et al., 2024; Liu et al., 2021; Somer et al., 2020). To our knowledge, only a few studies have combined rhythmic VS, oscillatory TES, and the online measurement of neural responses. SSVEPs were recovered during frequency-matched TES (Haslacher et al., 2021) or even enhanced by it (Dowsett et al., 2020; Ruhnau et al., 2016), but only lower frequencies up to 11 Hz have been tested. These reports suggest that modulating gamma-band SSVEPs through frequency-matched TES is a timely and important subject of investigation, but measurement feasibility and interaction effects are unclear.

The present study addresses this knowledge gap. We investigated A) if obtaining clean EEG data during gamma-band pulsed transcranial direct current stimulation, a form of oscillatory TES, is feasible; B) if gamma TES can augment SSVEPs elicited by gamma VS; and C) if different VS frequencies and TES sites influence the outcome. We tackled these questions in three experiments: In the first, VS and TES frequencies were closely matched around 40 Hz, and TES sites varied between occipito-central, centro-occipital (reversed polarity), and centro-frontal. Here, we aimed to rule out any effects of electrical currents on the retina or peripheral nerves as confounds (Asamoah et al., 2019; Kar & Krekelberg, 2012) and investigate potential impacts of current direction (Balslev et al., 2007). In the second experiment, occipito-central TES was applied just below 40 Hz, and VS frequency was either 35, 40, or 45 Hz. Here, we tested for frequency specificity. The third experiment included the same combined VS and TES conditions as the first experiment as well as control conditions where TES was on but VS was prevented from reaching the eyes by a blindfold. It served to quantify how effectively the data processing pipeline can mitigate electrical artefacts and separate them from real evoked brain activity.

To summarise, we hypothesised that it would be possible to record visually evoked 40 Hz activity with EEG during electrical brain stimulation. We also argued based on previous research that electrical and visual stimulation might interact, which could be specific to stimulation area (Experiment 1) and stimulation frequency (Experiment 2). Experiment 3 served as a control experiment, where we expected that under electrical brain stimulation, brain activity would be stronger and most similar to baseline SSVEPs when visual stimulation was also applied. Looking ahead, we were indeed able to successfully record 40 Hz SSVEPs with EEG during brain stimulation. Testing different sites and frequencies we did not find evidence for interactions.

## Materials & Methods

### Design

We hypothesised that to modulate visually evoked gamma activity, concurrent TES should be applied between occipital and central sites to target visual brain areas, and the frequency of both stimulation techniques should be closely matched (Dowsett et al., 2020; Ruhnau et al., 2016). Each factor (site and frequency) was tested in a different experiment. Experiments 1 and 2 included three blocks each: within every block, one trial of combined TES and VS was preceded and followed by a baseline trial with only VS at the same VS frequency (Figure 1A). In Experiment 1, VS was set to 40 Hz and TES to 39.9 Hz in all trials (see *Stimulation*), and TES sites varied per block. As depicted in Figure 1B, we selected anodal O2 to cathodal Cz (O2-Cz + or “occipital-central”) for the main condition, anodal Cz to cathodal O2 (Cz-O2 + “centro-occipital”) as a reversed polarity condition, and anodal Cz to cathodal Fz (Cz-Fz+ or “centro-frontal”) as a site control (Herring et al., 2019). In Experiment 2, 39.9 Hz TES was always anodal at O2 and cathodal at Cz, and VS frequency varied by block. The 40 Hz VS condition was equivalent to the occipito-central condition in Experiment 1; 35 and 45 Hz were frequency controls. The data from Experiments 1 and 2 revealed occasional residual TES artefacts appearing as square-shaped or spiked patterns, which could distort our results. Simulations revealed that this could be particularly problematic for the interpretation of Experiment 2, as residual artefacts in consecutive data segments time-locked to VS would be more likely to average out if VS and TES frequencies do not match (Supplementary Materials, Figure S1C). This prompted us to run another experiment to systematically optimise the artefact removal pipeline and objectively quantify its performance. Experiment 3 comprised two blocks (Figure 1A): The experimental block started with a baseline trial of only 40 Hz VS, followed by the same three TES conditions as in Experiment 1 - occipito-central, centro-occipital, and centro-frontal - with simultaneous 40 Hz VS. The defining feature of Experiment 3 was the “blackout” control block, where the VS device was on at 40 Hz, but the LEDs were covered with black tape. This precluded any visual stimulation effects while keeping control conditions as electrically equivalent as possible. The control block started with a resting state trial, with blackout VS and no TES. Three trials followed, matching the TES configurations of the experimental block but with blackout VS (Figure 1A). Any rhythmic signals captured in these trials could only result from residual TES artefacts, not SSVEPs, which cannot be evoked if VS light does not reach the retina.

**Figure 1:**
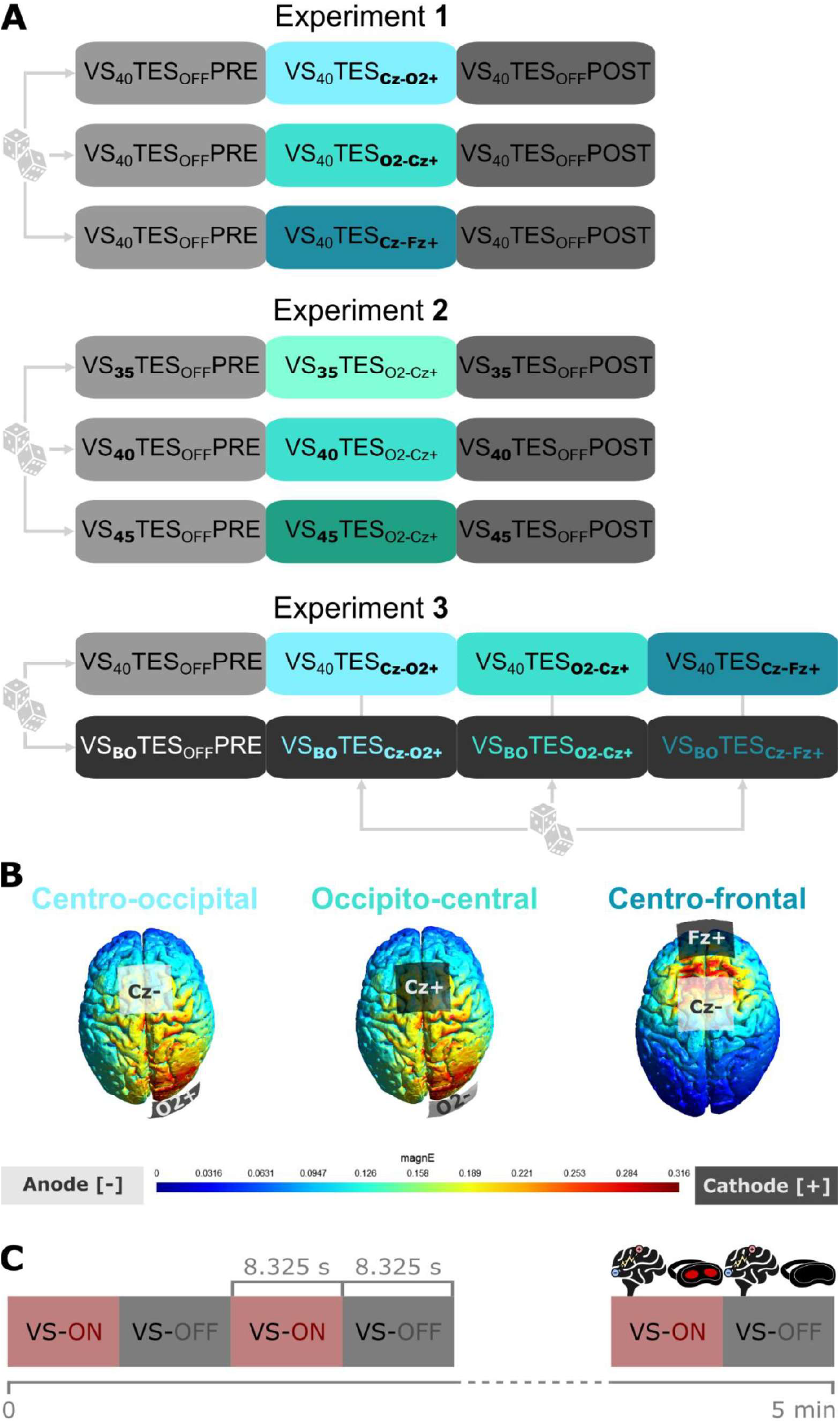
Experimental design. **A:** Levels of the independent variable manipulated within each experiment are in bold. Dice icons represent pseudorandomisation. VS = Visual Stimulation at a frequency of 35 Hz, 40 Hz, or 45 Hz, indicated by subscripts. In VS_BO_ (BO for “blackout”), the LEDs flickering at 40 Hz were covered with black tape. PRE = a trial without TES, recorded *before* a trial with TES; POST = a trial without TES, recorded *after* a trial with TES. TES = Transcranial Electrical Stimulation, either Cz anodal and O2 cathodal (Cz-O2+), O2 anodal and Cz cathodal (O2-Cz+), Cz anodal and Fz cathodal (Cz-Fz+) or no electrical stimulation (OFF), indicated by subscripts. **B:** Simulation of current flows for the three TES montages. The centro-occipital montage is equivalent to the occipito-central one with reversed polarity; they are equivalent here because the models display the absolute amplitude of the electric field. magnE = absolute magnitude of the electric field in V/m. **C:** Structure of all trials where TES was applied. TES was always on throughout the full 5 minutes of the trial. VS (blackout or visible) alternated between on for 8.325 seconds and off for 8.325 seconds.

### Protocol

Each session began with TES and EEG setup (impedances < 10 kOhm) and gradual habituation to TES and VS. There were three pseudorandomised blocks with three trials each in Experiments 1 and 2 and two pseudorandomised blocks with four trials each in Experiment 3. Here, the order of TES trials was also pseudorandomised and kept constant in both blocks within participants (Figure 1A). All trials were five minutes long. During trials, participants were instructed to keep their eyes closed and centred and to remain still. They took short breaks between blocks with the VS mask lifted, room lights on, and eyes open.

### Sample

With approval by the LMU Munich Psychology ethical committee and written informed consent obtained from all participants, we recruited 25 healthy adult volunteers per experiment. Ten volunteers took part in Experiments 1 and 2, three volunteers took part in Experiments 2 and 3, and eleven participated in all three Experiments. Their sessions were at least one month apart. The exclusion criteria were a history of seizures or epilepsy (also for first-degree relatives), any psychiatric or neurological condition, and colour blindness assessed with an Ishihara test. The demographic information is summarised in Table 1.

**Table 1:** Sample demographics. F=female; M=male; RH=right-handed; LH=left-handed; AD=ambidextrous. *All participants indicated that their gender matched their sex assigned at birth.

| Experiment | Sample size | Age mean (years) | Age range (years) | Sex* | Handedness |
| --- | --- | --- | --- | --- | --- |
| 1 | 25 | 23.08 | 18 - 28 | 18 F / 7 M | 21 RH / 3 LH / 1 AD |
| 2 | 25 | 23.76 | 18 - 28 | 17 F / 8 M | 21 RH / 3 LH / 1 AD |
| 3 | 25 | 26.4 | 18 - 30 | 15 F / 10 M | 22 RH / 3 LH |

### Stimulation

Transcranial Electrical Stimulation (TES) was administered using the neuroConn DC-Stimulator Plus (neuroCare Group GmbH, Munich, Germany). We used 5×5 cm square rubber electrodes with Ten20 conductive paste (Weaver and Company, Colorado, USA). The O2, Cz, and Fz positions were defined according to the 10-20 system, based on a previous study (Dowsett et al., 2020). Current flows simulated for a tDCS pulse using one author’s brain scans and simNIBS software (Saturnino et al., 2019) are shown in Figure 1B. The stimulation waveform was delivered to the stimulator with a digital-to-analogue converter using the “remote input” function. The pulsed transcranial direct current stimulation oscillated between 0 and 0.8 mA in a square-wave pattern with a duty cycle of 50 %. A squared pattern is more likely to modulate neuronal responses (Fröhlich & McCormick, 2010; Sherfey et al., 2018) and facilitates artefact removal (Dowsett et al., 2020); direct current allows for a test of polarity effects, as opposed to alternating current. The current strength of 0.8 mA was chosen as a compromise between participant comfort, artefact size, and effectiveness. The TES frequency of 39.9 Hz, just below the VS frequency of interest at 40 Hz, allowed for phase shifts between TES and VS, such that the VS and TES gradually drifted out of phase. We stimulated across all phases as the optimal stimulation phase is unknown; it may differ across participants, and multiple phases may elicit an effect. Cycling through all phases allows us to measure the average effect (Dowsett et al., 2020). Moreover, any TES artefacts remaining after data processing would be distributed across phase bins, averaging to zero given enough trials. During TES trials, TES was administered continuously for five minutes.

Visual stimulation (VS) was delivered through a custom-made system powered by a microcontroller, with LEDs embedded in a mask blocking off any external light. Stimulation (“flicker”) was temporally modulated in a square -wave pattern at a duty cycle of 50 %. Participants’ eyes were closed to match other experimental setups in our lab during sleep (Hainke et al., 2024). Narrowband red light with a peak wavelength of 605 nm facilitated transmittance through closed eyelids (Bierman et al., 2011); the target illuminance was 175 photopic lux at the eye, a value chosen based on pilot tests. In every trial, the temporally modulated light alternated between ON and OFF periods, each 8.325 s long (Figure 1C). This duration ensured that for the primary frequency of interest of 40 Hz, the TES and VS cycles drifted apart for long enough to cover all relative phases in one ON-period. Phase distributions were equally unbiased for 35 Hz and 45 Hz VS (Supplementary Materials, Figure S2). ON-periods originally containing both SSVEPs and TES artefacts were processed and then used for statistical analyses. OFF-periods containing only TES artefacts and no neural signal of interest were used for artefact removal. This applies equally to blackout trials in Experiment 3, where the LEDs were covered with black tape.

### EEG Acquisition & Processing

EEG data were recorded with a BrainAmp system (Brain Products GmbH, Gilching, Germany) at a 5 kHz sampling rate. The ground was placed on the left earlobe, the reference on the right earlobe, and the active electrode at the midpoint between the two main TES electrodes at O2 and Cz (near P2), where the electrical artefact would be smallest. Due to individual differences in head morphology, when needed, the active electrode was adjusted slightly until the raw data showed the characteristic TES square shape and did not clip. A DC correction was applied before every block to prevent signal saturation by the TES electric charge. Data were divided into short segments time-locked to the flicker, with a length depending on VS frequency of 22.2 ms (45 Hz), 25 ms (40 Hz), or 28.8 ms (35 Hz). Data were not filtered to prevent distorting the shape of the TES artefact.

To clean EEG data segments of TES artefacts, we developed an adaptive template subtraction method based on previous work by Dowsett et al. (2020). This pipeline and its parameters were optimised solely based on data from VS_BO_TES_O2-Cz+_, the control condition directly matching the main experimental condition included in all three experiments, VS_40_TES_O2-Cz+_. In VS_BO_TES_O2-Cz+,_ no VS was visible, so no neural signal of interest (i.e., SSVEP) could be evoked; also, segments were time-locked to the (unseen, blacked-out) 40 Hz VS and not to TES at 39.9 Hz. Thus, if the pipeline correctly removes TES artefacts, the segment average should be a relatively flat line. Any slight fluctuations should only reflect natural EEG noise, not a periodic 39.9 Hz signal. Simulations revealed how an average signal resembling a 40 Hz SSVEP could appear in control data if the artefact removal were to perform suboptimally in a larger number of consecutive segments (Supplementary Materials, Figure S1C). Therefore, modifications to the pipeline were deemed successful if they further minimised median peak-to-peak amplitudes of participant-level segment averages in the condition VS_BO_TES_O2-Cz+_. Only after the pipeline optimisation was concluded did we apply it universally to all data across all conditions and experiments, to prevent any bias of our statistical analyses.

Each segment-to-clean in VS-ON periods of TES trials underwent the processing pipeline visualised in Figure 2A and described as follows. First, the steepest points of the segment-to-clean were defined as the data points where its absolute first differential exceeded 10 % of the segment’s total amplitude (“steepest points”). The first differential of a time series quantifies the steepness of transitions between neighbouring time points. For a 25 ms segment, for example, this typically resulted in one subset of data points for the TES square-wave rise and one subset for the TES square-wave fall. These steep point subsets were each extended by two data points before and four data points after, to capture the full extent of the artefact (“artefact points’’). Then, segments potentially suitable for a template were selected from two VS-OFF periods - the one before and the one after the current VS-ON period (except if the segment-to-clean was in a trial’s first VS-ON period, then only the directly following VS-OFF period was available). VS-OFF data were digitally segmented with a sliding window matching the length of the segment-to-clean and moving forward one data point at a time. The peak-to-peak amplitude was computed for each VS-OFF segment minus the segment-to-clean; if this value was below 10 % of the segment-to-clean’s amplitude, the VS -OFF segment was pre-selected as a potential candidate for the template. Each pre-selected VS-OFF segment was assigned a score computed by subtracting the segment-to-clean from the VS-OFF segment, taking the first differential of this difference, and calculating the sum across the “artefact points”. The best combination of VS-OFF segments for a template was defined as the pair for which the absolute average score was the lowest. Because the first differential quantifies steepness between neighbouring data points, a low absolute score derived from the points most affected by the rising or falling artefact indicates a good match between segment-to-clean and template. The two best VS-OFF segments of approximately 300 were averaged into a template; then, the template was scaled to best match the amplitude of the segment-to-clean. The template was progressively scaled larger or smaller as long as this reduced the maximum of the absolute first differential at the “artefact points” of the segment-to-clean minus the scaled template. Lastly, the final template was baseline-corrected and subtracted from the baseline-corrected segment-to-clean.

**Figure 2:**
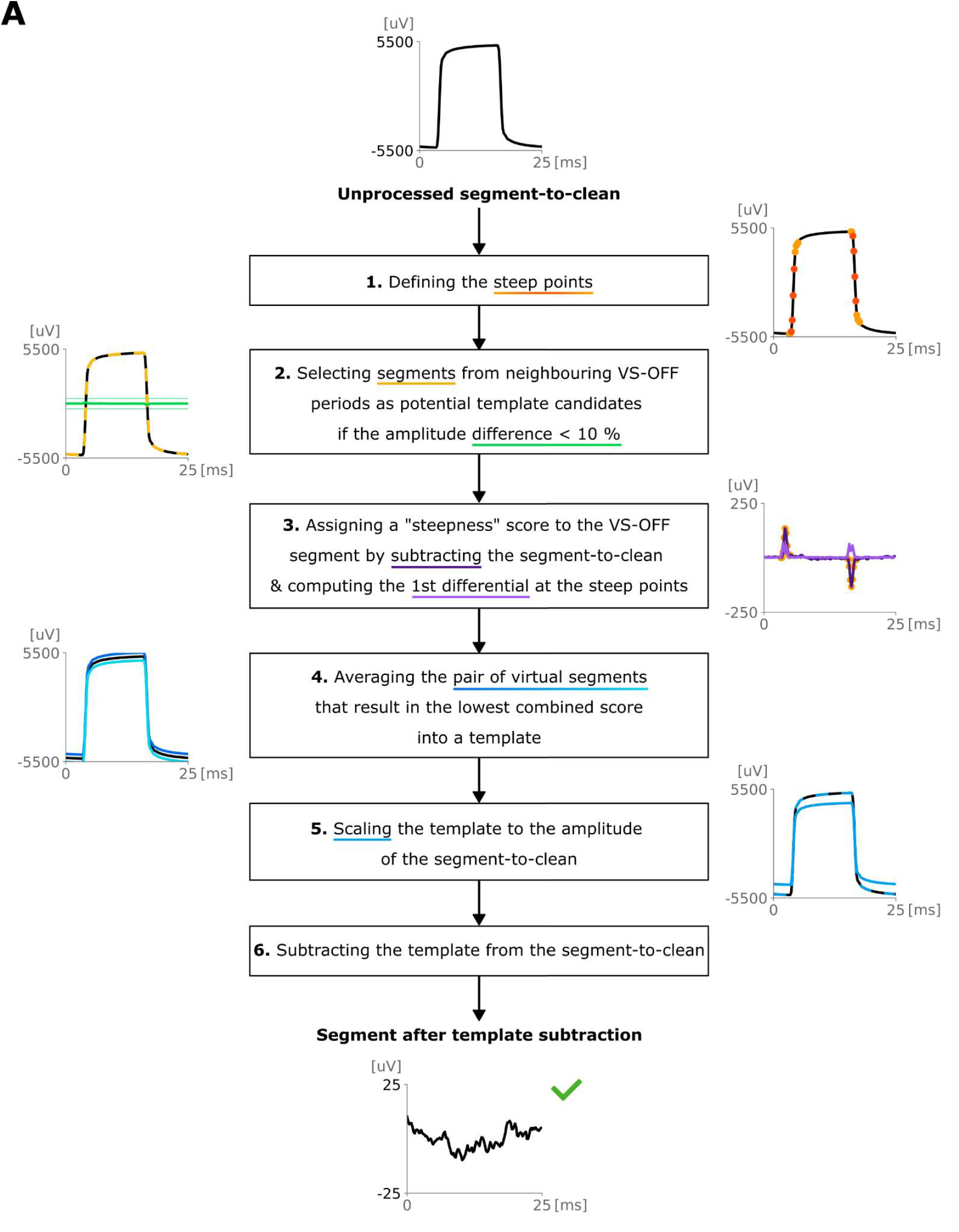
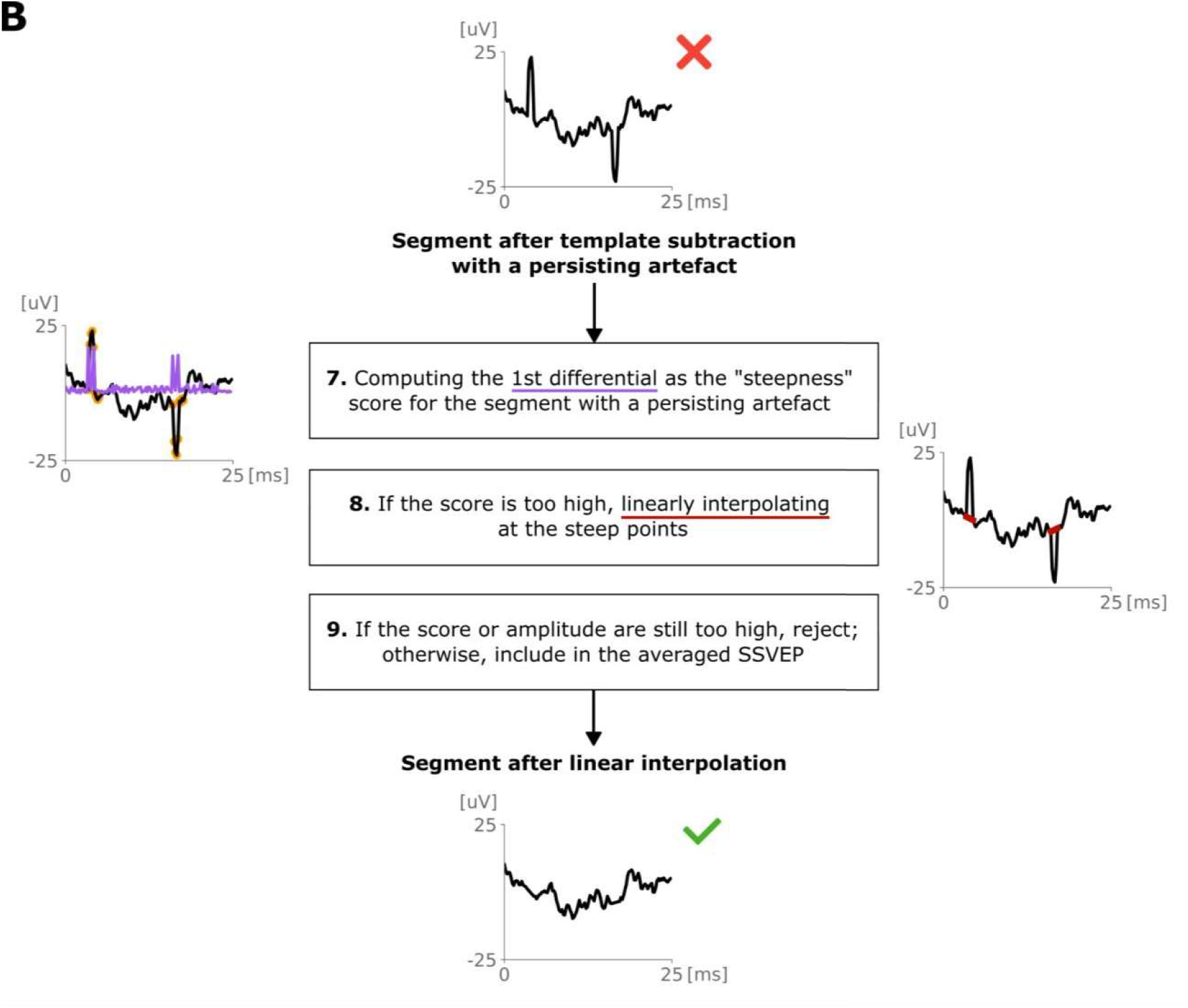
TES artefact removal pipeline. **A:** Artefact removal process for all segments in VS-ON periods, recorded during concurrent TES and time-locked to the flicker. Segment lengths of 25 ms depicted here correspond to 40 Hz VS, but the same applies to 22.2 ms or 28.8 ms segments (35 Hz or 45 Hz VS). **B:** Additional processing steps to mitigate artefacts surviving the processing pipeline depicted in A.

The next step of the pipeline, depicted in Figure 2B, dealt with any artefacts that resisted the above-mentioned procedure. They could either appear as patterns resembling the TES pulse square shape or as sharp spikes at the TES rise or fall points, in case the template’s shape or phase did not match the segment-to-clean well enough. This could result from a temporary increase of noise in the data, especially from a head movement. We first targeted the spike artefacts through linear interpolation: each segment after template subtraction was assigned a score, defined as the sum of its absolute first differential at the “steepest points”. If this score exceeded the systematically calibrated threshold of 16 pV, the “steepest points” subsets were each replaced with a straight line. If the segment’s score, recalculated after linear interpolation, still exceeded the threshold, a square-shaped artefact was likely present, so the segment was rejected.

Apart from these scores, three other criteria were defined for segment rejection post-processing: Segments in the first four seconds of each TES trial were discarded by default because muscular activity tended to be slightly higher at trial start, increasing the likelihood of residual artefacts. Second, segments for which less than two suitable template segments were found were removed. Lastly, as is common in standard EEG processing, segments with a peak-to-peak amplitude larger than 90 µV (likely to contain muscular artefacts) were excluded. This last step also applied to segments from TES_OFF_ conditions - which, free from TES artefacts by design, did not require any further processing. SSVEPs were finally computed by averaging all included segments by condition and participant into a time-domain Event-Related Potential (Supplementary Materials, Figures S3-S8).

### Statistical Analysis

In Experiments 1 and 2, we hypothesised an increase of SSVEP amplitudes only by frequency-matched TES applied between occipital and central sites. After-effects were also tested for in these conditions. In Experiment 3, we tested for residual TES artefacts by comparing TES-only conditions to resting-state data. Moreover, we hypothesised that SSVEPs evoked by 40 Hz VS could be recovered during frequency-matched TES at all sites, evidenced by A) larger amplitudes than the averaged signal from corresponding TES-only control data without VS and B) higher waveform correlations with baseline SSVEPs from VS-only data than with averaged signals from TES-only data. With a significance criterion of *a*=0.05 and Cohen’s *d* as effect size, directed hypotheses were tested through one-tailed permutation tests, and two-tailed permutation tests were run for expected null effects. Pearson correlation coefficients to assess SSVEP waveform similarity between two conditions were also subjected to permutation tests. Participants were excluded from a test if less than half of the data was available for any condition in that test.

## Results

The results of Experiment 1 are detailed in Table 2. Contrary to expectations, centro-occipital frequency-matched TES did not modulate SSVEP amplitudes during stimulation (H1.1), and occipito-central frequency-matched TES slightly decreased them (H1.3). Neither of these TES configurations induced after-effects on SSVEPs (H1.2, H1.4). As expected, centro-frontal frequency-matched TES did not increase SSVEP amplitudes during or after stimulation (H1.5, H1.6).

**Table 2:** Results of Experiment 1. Means, standard deviations (SD), and confidence intervals in microvolts. The symbol “<” indicates the direction of one-tailed tests; “=/=” represents two-tailed tests for expected null effects. PRE and POST refer to the VS-only trials before and after the TES condition within a given block, respectively – i.e., Cz-O2+ in H1.1 and H1.2, O2-Cz+ in H1.3 and H1.4, and Cz-Fz+ in H1.5 and H1.6.

| Variable 1 | Test | Variable 2 | N | Mean (SD) 1 | Mean (SD) 2 | Mean Difference & Confidence Interval | P-value | Cohen's d |
| --- | --- | --- | --- | --- | --- | --- | --- | --- |
| H1.1 $VS_{40}TES_{OFFPRE}$ | < | $VS_{40}TES_{Cz-O2+}$ | 25 | 1.44 (0.8) | 1.47 (0.75) | 0.04 [-0.11 ; 0.18] | 0.31 | 0.03 |
| H1.2 $VS_{40}TES_{OFFPRE}$ | < | $VS_{40}TES_{OFFPOST}$ | 25 | 1.44 (0.8) | 1.39 (0.84) | -0.05 [-0.15 ; 0.05] | 0.18 | -0.04 |
| H1.3 $VS_{40}TES_{OFFPRE}$ | < | $VS_{40}TES_{O2-Cz+}$ | 25 | 1.52 (0.83) | 1.36 (0.71) | -0.16 [-0.32 ; 0.01] | 0.03 | -0.14 |
| H1.4 $VS_{40}TES_{OFFPRE}$ | < | $VS_{40}TES_{OFFPOST}$ | 25 | 1.52 (0.83) | 1.48 (0.85) | -0.03 [-0.2 ; 0.13] | 0.32 | -0.03 |
| H1.5 $VS_{40}TES_{OFFPRE}$ | ≠ | $VS_{40}TES_{Cz-Fz+}$ | 25 | 1.46 (0.89) | 1.17 (0.6) | -0.29 [-0.58 ; -0.01] | 0.05 | -0.27 |
| H1.6 $VS_{40}TES_{OFFPRE}$ | ≠ | $VS_{40}TES_{OFFPOST}$ | 25 | 1.46 (0.89) | 1.39 (0.82) | -0.08 [-0.22 ; 0.07] | 0.27 | -0.06 |

The results of Experiment 2 are listed in Table 3. Frequency-matched occipito-central TES again did not increase SSVEP amplitudes during (H2.3) and after (H2.4) stimulation. As predicted, when VS frequency was lower (35 Hz) or higher (45 Hz) than TES frequency, occipito-central TES did not modulate SSVEP amplitudes during stimulation (H2.1, H2.5). SSVEP amplitudes after TES were unaltered in the 35 Hz VS condition (H2.2) and decreased in the 45 Hz condition (H2.6).

**Table 3:**
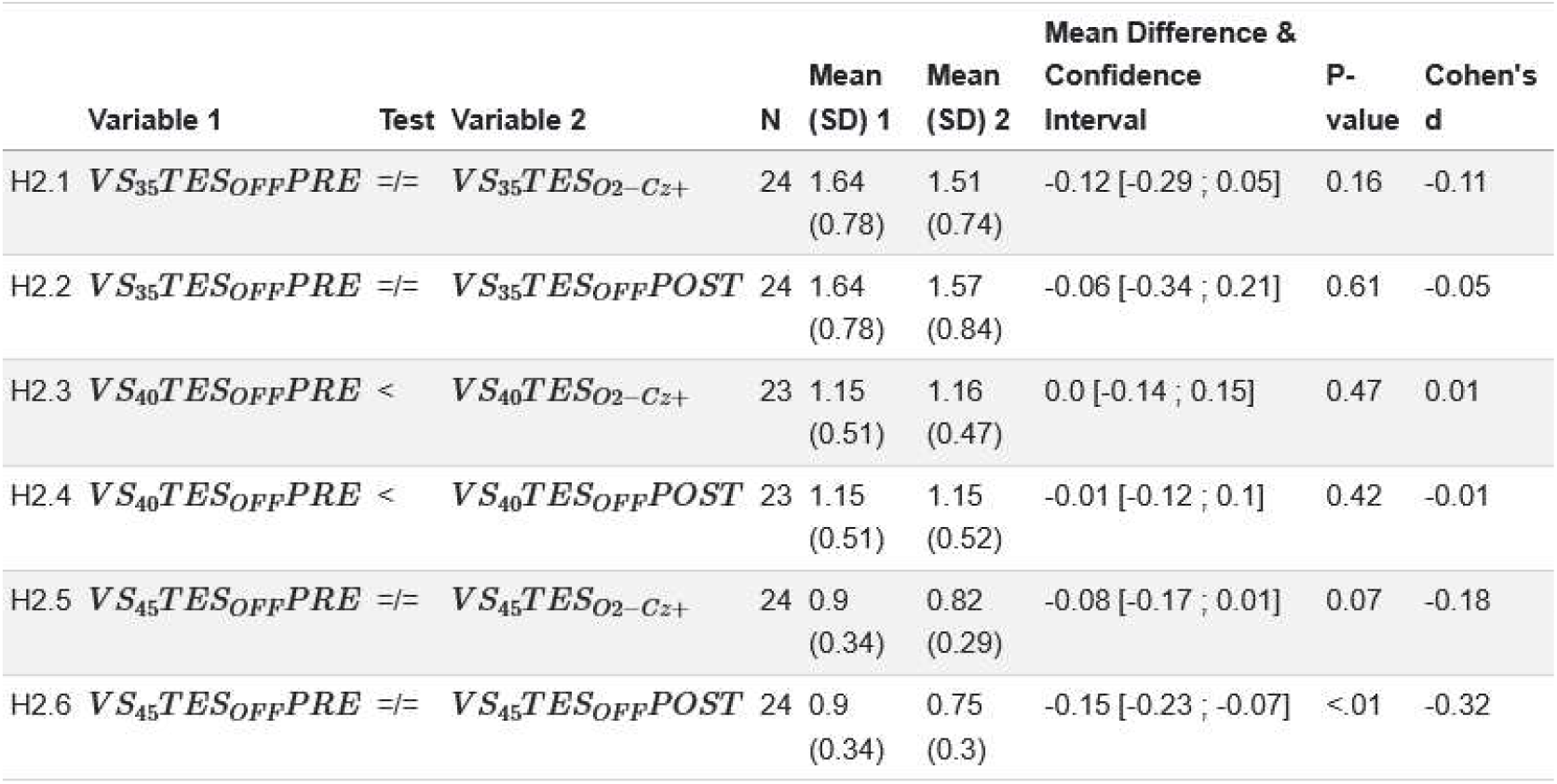
Results of Experiment 2. Means, standard deviations (SD), and confidence intervals in microvolt. The symbol “<” indicates the direction of one-tailed tests; “=/=” represents two-tailed tests for expected null effects.

| Variable 1 | Test | Variable 2 | N | Mean (SD) 1 | Mean (SD) 2 | Mean Difference & Confidence Interval | P-value | Cohen's d |
| --- | --- | --- | --- | --- | --- | --- | --- | --- |
| H2.1 $VS_{35}TES_{OFFPRE}$ | ≠ | $VS_{35}TES_{O2-Cz+}$ | 24 | 1.64 (0.78) | 1.51 (0.74) | -0.12 [-0.29 ; 0.05] | 0.16 | -0.11 |
| H2.2 $VS_{35}TES_{OFFPRE}$ | ≠ | $VS_{35}TES_{OFFPOST}$ | 24 | 1.64 (0.78) | 1.57 (0.84) | -0.06 [-0.34 ; 0.21] | 0.61 | -0.05 |
| H2.3 $VS_{40}TES_{OFFPRE}$ | < | $VS_{40}TES_{O2-Cz+}$ | 23 | 1.15 (0.51) | 1.16 (0.47) | 0.0 [-0.14 ; 0.15] | 0.47 | 0.01 |
| H2.4 $VS_{40}TES_{OFFPRE}$ | < | $VS_{40}TES_{OFFPOST}$ | 23 | 1.15 (0.51) | 1.15 (0.52) | -0.01 [-0.12 ; 0.1] | 0.42 | -0.01 |
| H2.5 $VS_{45}TES_{OFFPRE}$ | ≠ | $VS_{45}TES_{O2-Cz+}$ | 24 | 0.9 (0.34) | 0.82 (0.29) | -0.08 [-0.17 ; 0.01] | 0.07 | -0.18 |
| H2.6 $VS_{45}TES_{OFFPRE}$ | ≠ | $VS_{45}TES_{OFFPOST}$ | 24 | 0.9 (0.34) | 0.75 (0.3) | -0.15 [-0.23 ; -0.07] | <.01 | -0.32 |

Table 4 summarises the results of Experiment 3. While TES artefacts are unlikely to be eliminated entirely (Kasten & Herrmann, 2019; H3.1, H3.3, H3.5), here they have been successfully minimised to a level smaller than the neural signal of interest. The averaged signal from conditions with both visual and electrical stimulation, baseline-corrected by data unaffected by any stimulation, had a larger amplitude than in conditions with electrical stimulation alone, equally baseline-corrected. This is true for occipito-central and centro-occipital TES (H3.2, H3.4); the test for centro-frontal TES did not reach significance (H3.6). It follows that apart from centro-frontal TES conditions, recording neuronal responses evoked by VS with EEG was possible despite the ongoing brain stimulation with occipital and central TES (Figure 3).

**Table 4:** Results of Experiment 3. Means, standard deviations (SD), and confidence intervals in microvolts, except for H3.7 and H3.8 (unit-free correlation coefficients). The symbol “<” indicates the direction of one-tailed tests; “=/=” represents two-tailed tests for expected null effects; “X” stands for Pearson correlations of SSVEP waveforms.

| Variable 1 | Test | Variable 2 | N | Mean (SD) 1 | Mean (SD) 2 | Mean Difference & Confidence Interval | P-value | Cohen's d |
| --- | --- | --- | --- | --- | --- | --- | --- | --- |
| H3.1 $VS_{BO}TES_{OFFPRE}$ | ≠ | $VS_{BO}TES_{Cz-O2+}$ | 21 | 0.32 (0.07) | 0.66 (0.21) | 0.34 [0.24 ; 0.45] | <.01 | 1.56 |
| H3.2 $VS_{BO}TES_{Cz-O2+} - VS_{BO}TES_{OFFPRE}$ | < | $VS_{40}TES_{Cz-O2+} - VS_{BO}TES_{OFFPRE}$ | 21 | 0.34 (0.23) | 0.72 (0.42) | 0.37 [0.18 ; 0.56] | <.01 | 0.78 |
| H3.3 $VS_{BO}TES_{OFFPRE}$ | ≠ | $VS_{BO}TES_{O2-Cz+}$ | 20 | 0.31 (0.07) | 0.67 (0.23) | 0.35 [0.25 ; 0.45] | <.01 | 1.49 |
| H3.4 $VS_{BO}TES_{O2-Cz+} - VS_{BO}TES_{OFFPRE}$ | < | $VS_{40}TES_{O2-Cz+} - VS_{BO}TES_{OFFPRE}$ | 20 | 0.35 (0.21) | 0.93 (0.51) | 0.58 [0.33 ; 0.82] | <.01 | 1.04 |
| H3.5 $VS_{BO}TES_{OFFPRE}$ | ≠ | $VS_{BO}TES_{Cz-Fz+}$ | 18 | 0.31 (0.06) | 0.62 (0.2) | 0.3 [0.2 ; 0.41] | <.01 | 1.44 |
| H3.6 $VS_{BO}TES_{Cz-Fz+} - VS_{BO}TES_{OFFPRE}$ | < | $VS_{40}TES_{Cz-Fz+} - VS_{BO}TES_{OFFPRE}$ | 18 | 0.3 (0.21) | 0.41 (0.21) | 0.11 [-0.06 ; 0.27] | 0.09 | 0.36 |
| H3.7 $VS_{40}TES_{O2-Cz+} \times VS_{BO}TES_{O2-Cz+}$ | < | $VS_{40}TES_{O2-Cz+} \times VS_{40}TES_{OFFPRE}$ | 20 | 0.27 (0.54) | 0.7 (0.33) | 0.44 [0.12 ; 0.76] | <.01 | 0.7 |
| H3.8 $VS_{40}TES_{Cz-O2+} \times VS_{BO}TES_{Cz-O2+}$ | < | $VS_{40}TES_{Cz-O2+} \times VS_{40}TES_{OFFPRE}$ | 21 | -0.04 (0.52) | 0.76 (0.38) | 0.8 [0.51 ; 1.09] | <.01 | 1.25 |

**Figure 3:**
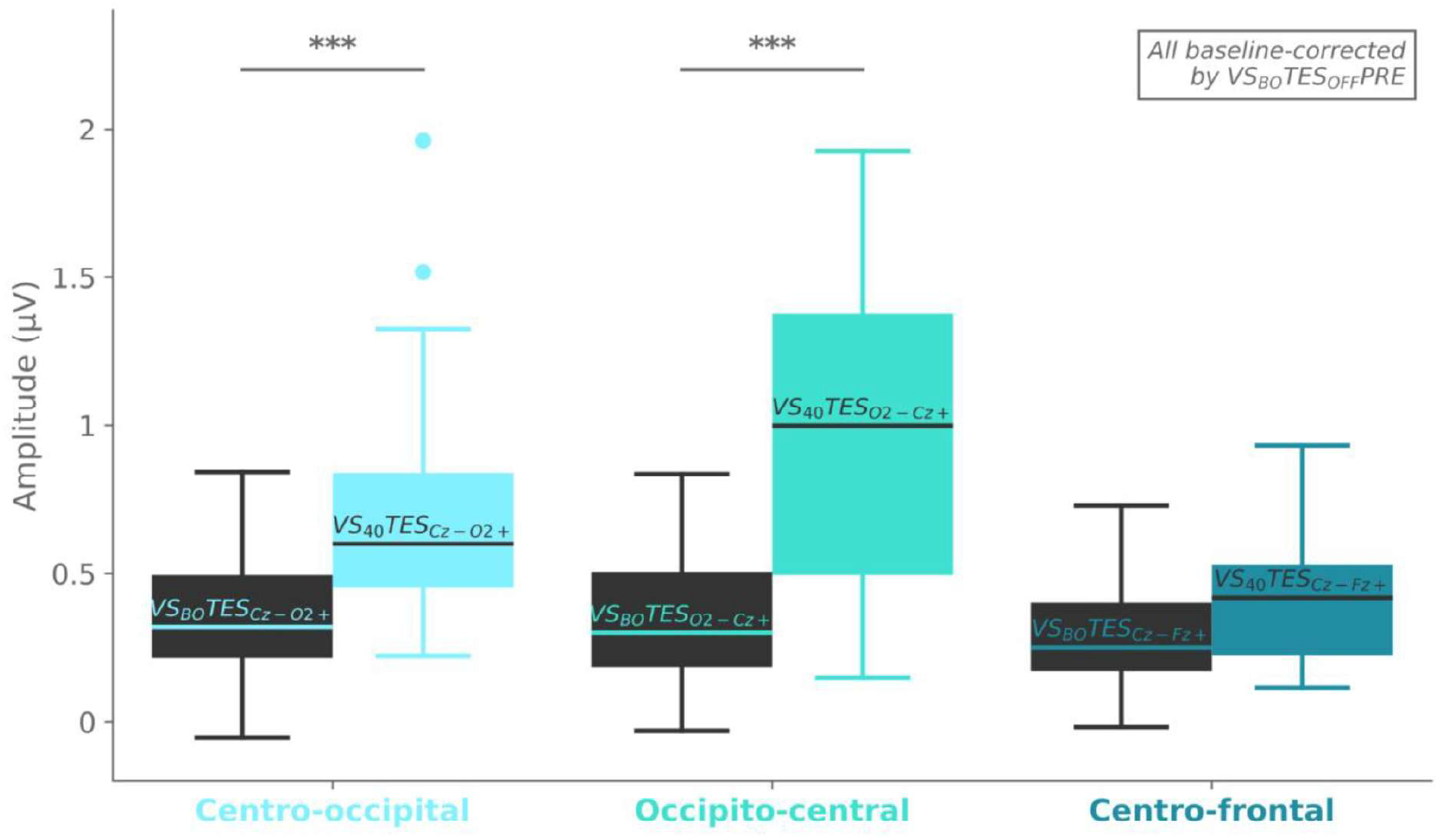
Visual responses (SSVEPs) successfully recovered with EEG during TES applied between central and occipital electrodes. Peak-to-peak amplitudes of averaged segments by condition in Experiment 3. Black boxplots represent conditions with blackout VS (VS_BO_) and active TES; these signals may contain residual TES artefacts, but no visually evoked neuronal activity. Coloured boxplots represent the equivalent conditions with visible 40 Hz VS (VS_40_); these signals should additionally contain visually evoked neuronal activity at 40 Hz. All were baseline-corrected by the resting-state condition with blackout VS and no TES (VS_BO_TES_OFF_PRE). Boxplot lines mark median values; boxes delimit the interquartile range; whiskers encompass data points within 1.5x of the interquartile range from box limits; ^***^ = *p*<.001.

SSVEP waveform correlations further supported this finding. Note that while TES-only trials may contain residual artefacts but no visually evoked neural signal, the opposite is true for VS-only trials. Our data show that the averaged signal from the conditions combining VS and TES was more highly correlated with the signal acquired during VS-only than with the signal acquired during TES-only (H3.7, H3.8). For a more qualitative inspection, Figure 4 displays the participant-level SSVEPs. Regardless of whether TES was applied simultaneously or not, recovered waveforms tended to be noticeably similar within participants, sinusoidal, and consistent with a VS frequency of 40 Hz in terms of period.

**Figure 4:**
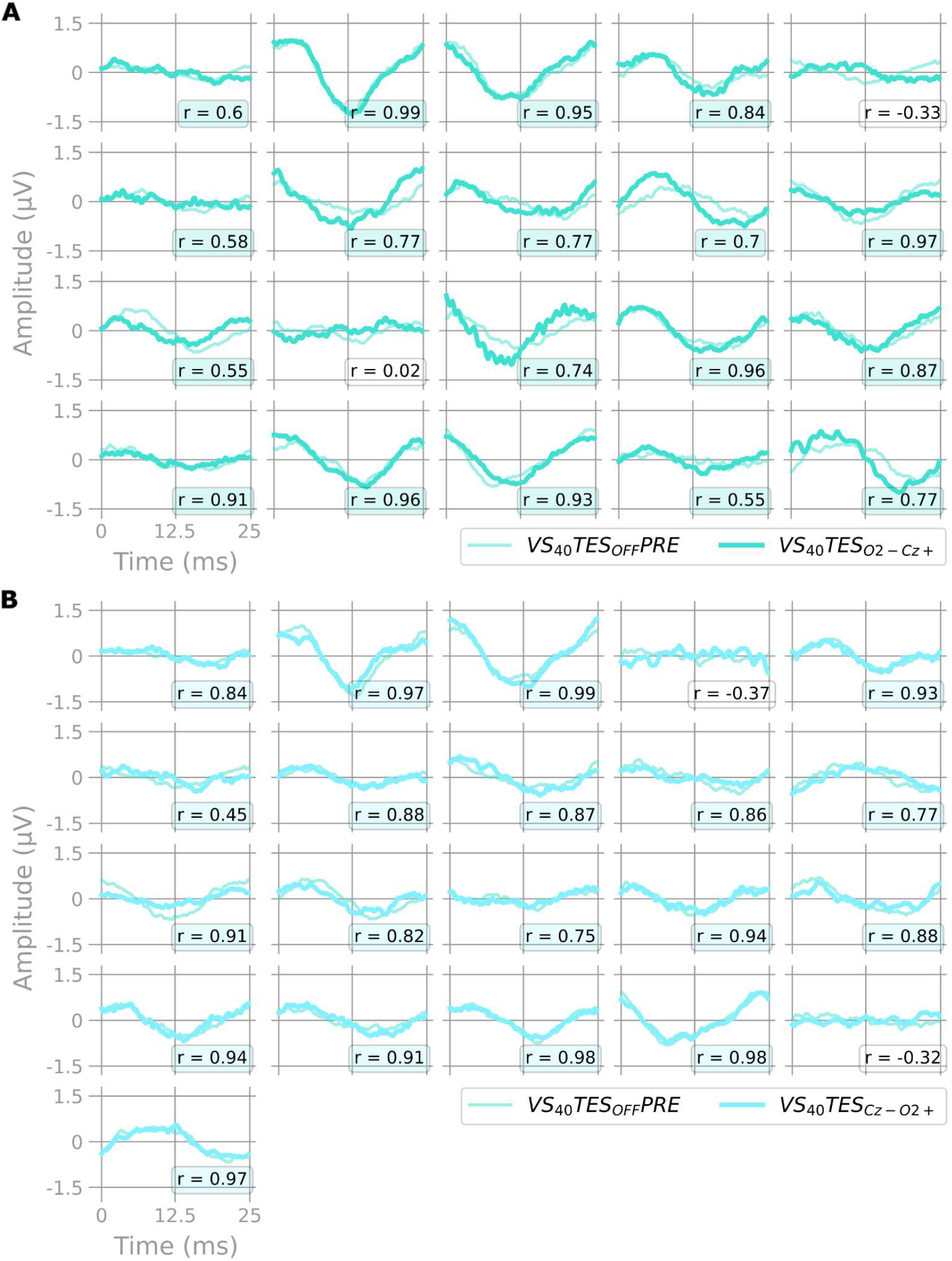
40 Hz SSVEPs, recorded with or without concurrent TES, were highly correlated within participants. SSVEPs in the time domain at the participant level. Data were taken from Experiment 3 and processed as described in *EEG Acquisition & Processing*. Within-participant correlation coefficients of the two conditions’ average waveforms are presented at the bottom right of every subplot and coloured if *r* > 0.4. **A:** SSVEPs from the occipito-central condition. VS_40_TES_OFF_PRE: 40 Hz Visual Stimulation alone. VS_40_TES_O2-Cz+_: 40 Hz Visual Stimulation and concurrent Transcranial Electrical Stimulation at 39.9 Hz, anodal at the occipital site O2 and cathodal at the central site Cz. **B:** SSVEPs from the centro-occipital condition. VS_40_TES_OFF_PRE: 40 Hz Visual Stimulation alone, the same as in A. VS_40_TES_Cz-O2+_: 40 Hz Visual Stimulation and concurrent Transcranial Electrical Stimulation at 39.9 Hz, anodal at the central site Cz and cathodal at the occipital site O2.

## Discussion

In sum, the present study reveals that recording brain responses to 40 Hz visual flicker during electrical brain stimulation is possible. Capturing neuronal signals with EEG during TES is a persistent challenge in the neuromodulation field because TES currents are much larger than neuronal signals; this challenge is further exacerbated when neuronal frequencies such as gamma are particularly low in amplitude and when TES is frequency-matched to VS. Here, we present a method that successfully recovers 40 Hz SSVEPs during frequency-matched oscillatory TES. Although our data do not suggest that frequency-matched TES augments 40 Hz SSVEPs, through signal amplitudes and waveforms, the data do strongly indicate that the recovered signal represents neuronal activity rather than electrical or physiological confounds. These findings open the door for fundamental and clinical studies to combine rhythmic sensory and electrical stimulation in the gamma band and concurrently verify neuronal effects with EEG.

To summarise Experiments 1 and 2, we tested if oscillating TES could increase SSVEP amplitudes, depending on electrical stimulation areas and the frequency overlap between TES and VS. Our data did not support this hypothesis, for which there may be several reasons. It may be that TES just below 40 Hz does not affect ongoing 40 Hz responses to visual flicker: visual responses are generated by parallel dynamic processing across billions of neurons, and TES may have been too coarse or weak to affect them. Strong visual responses such as those elicited by visual flicker, especially square-wave flicker (Panitz et al., 2023), could be particularly hard to disrupt. Two effects from Experiments 1 and 2 did reach significance, but given the recent controversy over the efficacy of TES (Parkin et al., 2015; Polanía et al., 2018), we are careful not to overinterpret them. For one, occipito-central TES appears to have reduced 40 Hz SSVEP amplitudes in Experiment 1. However, the effect size was low (*d* = −0.14), and the result did not reproduce in Experiments 2 (Results, H2.3) or 3 (*post*–*hoc: p* = .09, *d* = +0.19). For another, 45 Hz SSVEP amplitudes were lower after occipito-central TES just below 40 Hz in Experiment 2. This is likely to be a coincidental finding, given the low effect size (*d* = -0.32) and the absence of after-effects in all other conditions. Overall, fatigue due to the long experimental sessions may have introduced variability in the SSVEPs.

One possibility to consider is that the TES at 39.9 Hz drifting in and out of phase with 40 Hz VS may have enhanced the SSVEPs at some phases but inhibited them at the opposite phase. In this work, we prioritised the validation of the data processing pipeline; because the literature shows that fully eliminating TES artefacts remains challenging (Kasten & Herrmann, 2019), we reasoned that an equal distribution of TES-VS phases would allow for a more solid interpretation of results (Dowsett et al., 2020). Moreover, there is no clear indication in the literature yet as to what exact TES-VS phase would be most effective. Now that we have validated the artefact removal pipeline, future work could try to systematically vary the phase between VS and frequency-matched TES. It is conceivable that TES could enhance SSVEP amplitudes in a subset of phases, and that the optimal phase exhibits interindividual variability. The results would be reliably interpretable as long as recovered SSVEPs are larger than averaged signals in a control condition with VS at the same phase but blacked out. One more point to mention is the TES current strength of 0.8 mA, which we chose as a compromise between participant comfort, artefact size, and the likelihood of an effect. A higher current strength could have modulated SSVEP amplitudes but also increased participant discomfort. This could have led to muscular contraction, impairing artefact removal. Future studies could test higher current intensities by applying anaesthetic gels to the skin to reduce stimulation sensation (Kerstens et al., 2022). It would also be interesting to explore to what extent other TES protocols such as transcranial alternating current stimulation or sawtooth-shaped TES may modify the outcome (Dowsett et al., 2020; Dowsett & Herrmann, 2016).

As for Experiment 3, we tested how reliably we could recover 40 Hz SSVEPs despite large TES artefacts, depending on electrical stimulation areas. We confirmed that the signal during combined visual and electrical stimulation applied to occipital and central areas was larger than during electrical stimulation without additional visual stimulation (Figure 3). Although fully eliminating TES artefacts remains difficult (Kasten & Herrmann, 2019), the pipeline presented here achieves a reduction of TES artefact magnitude by a factor of 220: in the main control condition (occipito-central TES and blackout VS), for instance, segment amplitudes dropped from an average of 6356 μV in the raw data to an average of 28.9 μV after processing. This value is well within the range of 0.5−100µV common for resting-state EEG data (Teplan, 2002). Processing and then averaging short data segments allowed for a more fine-grained artefact template subtraction approach, which can handle variations in TES artefacts over time caused by changes in impedance levels, for example. Delimiting segments to the duration of one VS cycle resulted in a large number of segments (e.g., 6000 25-millisecond segments in 40 Hz VS trials) and consequently, a high signal-to-noise ratio: confounds such as muscular or ocular activity, which share the gamma frequency range (Hipp & Siegel, 2013) but are not in phase with VS, average out. Our results suggest that with enough segments, Event-Related Potentials are interpretable despite simultaneous TES, which opens up a range of possible applications.

Our approach works well for EEG electrodes placed between two TES electrodes; it has yet to be adapted for EEG electrode positions further away from the TES electrode midline. In the centro-frontal TES condition, the TES artefact was too large to be mitigated sufficiently and to avoid EEG signal saturation in all trials. This may have resulted from a large difference in the electric field at the active and reference EEG electrodes, due to their relative position to the TES electric field on the scalp. To tackle this issue, future studies could test alternative types of EEG electrodes, different TES electrode sizes, or other TES waveforms with sharp transitions like sawtooth waves. Nonetheless, the results of Experiment 3 uncovered how solid control conditions, in which TES is on but sensory stimulation is prevented from reaching the relevant sensory organ, are necessary to interpret data recovered during TES. This is particularly important when different stimulation frequencies are compared, given that residual electrical artefacts may have a higher impact on SSVEP outcome when TES and VS frequencies match (Supplementary Materials, Figure S1). Matching stimulation frequencies seems to be the most useful approach, though, considering the clear focus on 40 Hz throughout gamma stimulation literature (Blanco-Duque et al., 2023; Iaccarino et al., 2016). To strengthen our claim that we successfully recovered 40 Hz SSVEPs during TES, we inspected the SSVEP waveforms: Our data showed that SSVEPs recovered during centro-occipital and occipito-central TES were more similar to baseline SSVEPs, recorded without concurrent TES, than to the average signal from TES-only data, which may contain residual artefacts but no neural signal of interest. With or without TES applied simultaneously, SSVEPs showed sinusoidal waveforms and periods consistent with a VS frequency of 40 Hz. The similarity within participants is visible (Figure 4) and statistically significant.

Overall, the present work establishes an optimised method to clean high-frequency EEG data from periodic electrical artefacts, laying the ground for a range of applications combining the increasingly popular Rhythmic Sensory Stimulation and the widely used Transcranial Electrical Stimulation (Hanslmayr et al., 2019; Koch et al., 2024; Thut et al., 2011; D. Wang et al., 2024). Using the method presented, future work can systematically explore how frequency-matched TES might modulate sensory evoked potentials under different conditions. On the one hand, sensory and electrical stimulation could be matched at a different frequency of choice. Frequency bands below gamma, both endogenous and evoked, tend to be stronger due to the natural 1/f distribution of EEG frequency power (Herrmann, 2001). This might explain why our results differ from those of Dowsett et al. (2020), who stimulated in the alpha band. Moreover, stimulation could be adapted to peak individual frequencies, i.e., the frequency within a given range such as gamma for which an individual exhibits the largest evoked potential (Mockevičius et al., 2023). On the other hand, our technique could also be applied to other sensory domains, particularly auditory and tactile (Jones et al., 2020; Mosabbir et al., 2022; Rufener et al., 2023). This would extend the usability of our multimodal protocol to individuals with impaired eyesight and interventions targeting other sensory pathways (Clements-Cortes et al., 2016).

To conclude, we demonstrated that 40 Hz SSVEPs can be reliably measured during frequency-matched Transcranial Electrical Stimulation, providing evidence in favour of recovered neuronal activity distinct from electrical or physiological confounds. These findings lay the groundwork for quantifying neuronal responses to combined rhythmic sensory and electrical stimulation in the gamma band. Such a combination is likely to be clinically relevant for individuals with altered endogenous gamma oscillations: both sensory and electrical gamma stimulation protocols carry potential as adjunctive treatment options for Mild Cognitive Impairment and Alzheimer’s Disease (Shu et al., 2024; Traikapi & Konstantinou, 2021; H. Wang et al., 2025), Major Depressive Disorder (Fitzgerald & Watson, 2018), and Schizophrenia (Black et al., 2024; Palmisano et al., 2024). Whether or not the stimulation techniques may interact, the possibility of increasing clinical benefits through a multimodal intervention merits further investigation. Importantly, neuronal responses during stimulation should be quantified, as the modulation of neuronal gamma activity is deemed necessary for clinical effects. The technique presented here sets the stage for such endeavours.

## Abbreviations

EEG: Electroencephalography
SSVEP: Steady-State Visually Evoked Potential
TES: Transcranial Electric Stimulation
VS: Visual Stimulation

## Acknowledgements

The authors would like to acknowledge Joshi Walzog and Oliver Holder from the Max-Planck Institute for Biological Cybernetics electronics workshop for developing the visual stimulation system, and all participants.

## Data Availability

This study was preregistered on Open Science Framework (https://doi.org/10.17605/OSF.IO/JD3NQ) and preprinted on BioRXiv (https://doi.org/10.1101/2024.06.21.599984). Upon publication, all anonymised data will be shared on Open Science Framework. Study materials, code for data processing, and notebooks for statistical analyses will be made available on GitHub.

## Author Contributions

Conceptualisation: LH, JD, PT, MS

Data Curation: LH

Formal Analysis: LH, JD, PT, MS

Funding Acquisition: JD, PT, JP

Investigation: LH, JD

Methodology: LH, JD, PT

Project Administration: JD, PT

Resources: JD, PT

Software: LH, JD

Supervision: JD, PT, MS, JP

Validation: LH, JD

Visualisation: LH

Writing Original Draft Preparation: LH

Writing Review & Editing: JD, PT, MS, JP

## Funding

This work was supported by the Deutsche Forschungsgesellschaft (DFG) grants [DO 2460/1-1] to JD and [TA857/3-2] to PT, the Excellence Strategy of the Federal Government and the Länder through the NEUROTECH Innovation Network (IGSSE) to JP, and the Max Planck Society to MS.

## Declarations of Interest

LH, MS, PT, and JD report no conflicts of interest. JP received consulting fees from Uniqure, has a patent on Epo variants, is on the advisory board of the Sinapps2 study, and is a board member of the following societies: DGPPN, DGGPP, DZPG, DZNE.

## Notes

### Summary of Updates

Upon reinspection of the data from experiments 1 and 2, optimisating the artefact removal pipeline and objectively quantifying its performance was demeed necessary. Therefore, we conducted a third experiment. While the original results from experiments 1 and 2 had to be revised, experiment 3 substantiated the methodological advance presented in our updated manuscript.

