## Supplementary_materials for "40 Hz Steady-State Visually Evoked Potentials Recovered During Oscillating Transcranial Electrical Stimulation"

### Simulated impact of transient noise in EEG data during TES

Figure S1 depicts a simulation of how transient noise in the EEG data during TES can affect SSVEP outputs. An example of such noise could be brisk participant movement.
First, we simulated a square wave with an arbitrary amplitude of one and a period corresponding to 39.9 Hz to represent the TES shape in raw EEG data. This square wave was concatenated 333 times to simulate one full VS-ON period. The amplitude of the middle 20 % of square waves was doubled, simulating a short period of high noise in the data. Then, we added triggers at a rate of 40 Hz, marking segment lengths equivalent to conditions with 40 Hz VS. Figure S1A shows the entire simulated data; Figure S1B is the same plot but zoomed into a shorter chunk of data to visualise how the simulated 40 Hz VS triggers and the 39.9 Hz TES slowly drift apart.

We ran simulations for 35 Hz, 40 Hz, and 45 Hz VS triggers. The simulated TES data were always the same; only the trigger times differed. For visualisation, we computed segment averages stratified into five phase bins, representing relative phases of TES to VS trigger. In phase bin 1, the TES square wave rise was at or shortly after the VS trigger; this square wave rise progressively moved away from the VS trigger with increasing phase bin number. Figure S1C shows the segment averages by phase bins in colours and the average across all segments in black for each VS trigger frequency. While segments average to a flat line in the 35 Hz and 45 Hz simulations, the 40 Hz simulation shows a pattern that resembles a sinusoidal SSVEP in the time domain with a near-40 Hz period.

The most likely explanation is that TES artefact size distortion affecting multiple consecutive segments spreads across more phase bins when TES and VS trigger frequencies do not match closely, as was the case for simulated 35 Hz and 45 Hz VS triggers. Simulated 40 Hz VS triggers, however, only drifted apart from simulated 39.9 Hz TES slowly. The 40 Hz subplot of Figure S1C reveals that only the neighbouring bins 2 and 3 have a higher amplitude, skewing the overall segment average toward their shape. This simulation reveals the importance of inspecting time-series data acquired under simultaneous VS and TES after processing to ensure that noise sources have been dealt with appropriately. Particularly, it highlights that it is crucial for a solid experimental design to include control conditions without the sensory stimulation that should evoke the neuronal signal of interest (here, VS) but with active TES equivalent to the “real” experimental conditions. If the artefact removal pipeline could not handle temporary periods of noise in the EEG sufficiently, this would be revealed in the control conditions. By contrasting experimental conditions evoking the neuronal signal of interest with electrically equivalent control conditions, we are able to disentangle neuronal from artefactual sources of recovered periodic EEG signals.

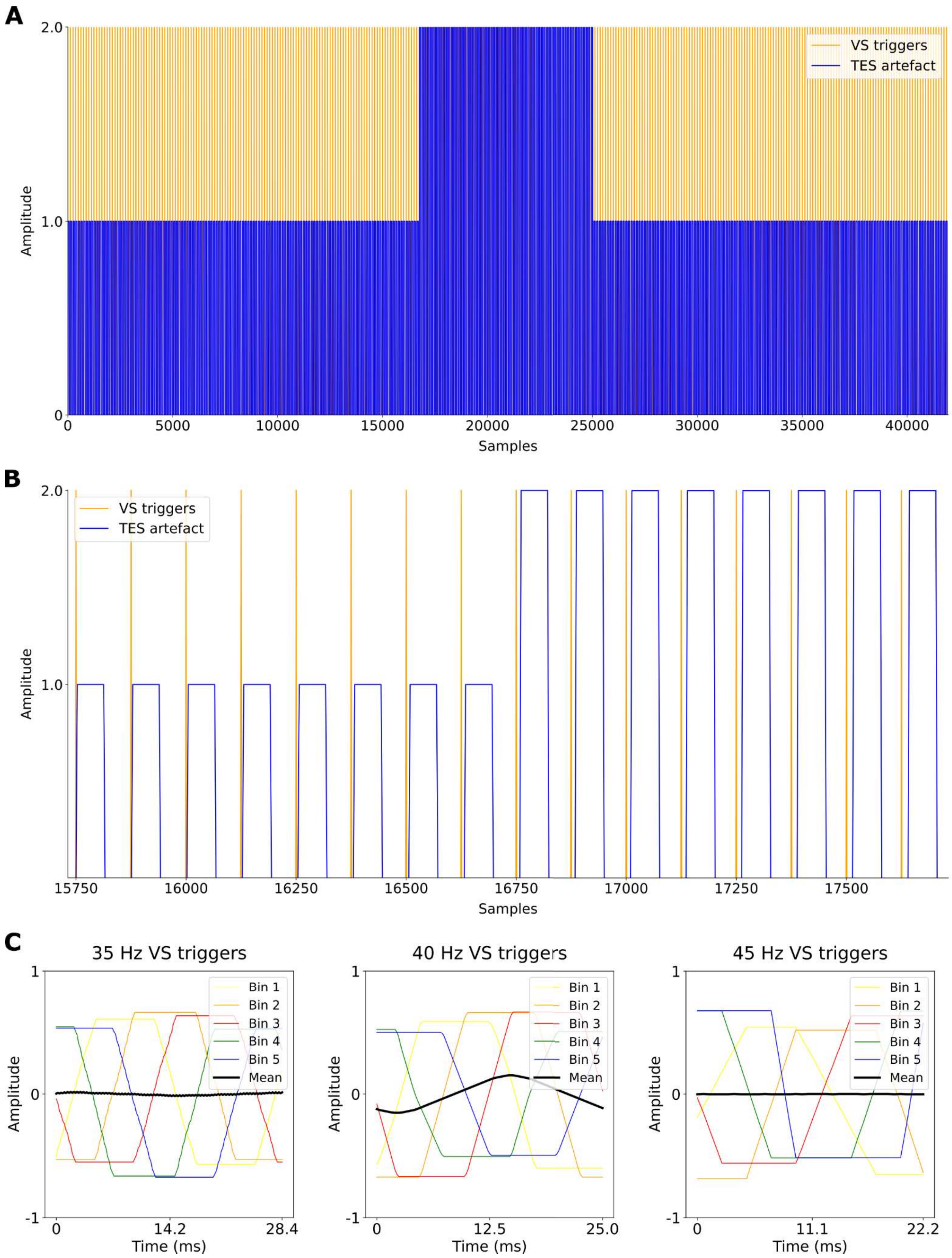


**Figure S1: Simulated increase in TES artefact size mid-trial affects segment averages depending on VS frequency.**

**A:** Simulation of square-shape 39.9 Hz TES artefacts and segment triggers corresponding to 40 Hz VS, for a VS-ON period of 8.325 seconds. In the middle 20 % of data, TES artefacts are twice as large, simulating an increase of noise in EEG data. **B:** The same plot as in A but zoomed in to visualize how the simulated VS and TES drift out of phase. **C:** Segment averages by phase bins (coloured) and overall mean (black), by simulated VS frequency.

### Phase distributions between TES and VS

In VS-ON periods of TES trials, all phases between 39.9 Hz TES and VS should be cycled through and no phase should be overrepresented, regardless of VS frequency (35 / 40 / 45 Hz). To verify this, we computed the phases by extracting the maximum absolute first differential for each VS-ON segment as the index for the steepest rise in the data, corresponding to the onset of square-wave TES. For 35 Hz, segments were longer than for 40 Hz, so only the first steep rise in each segment was extracted. For 45 Hz, segments were shorter than for 40 Hz, so segments were padded with 2 ms before and after to match a length of 25 ms equivalent to 40 Hz and rejected if the steep rise occurred outside segment boundaries. Indices were converted to radians; group average values are depicted in Figure S2.

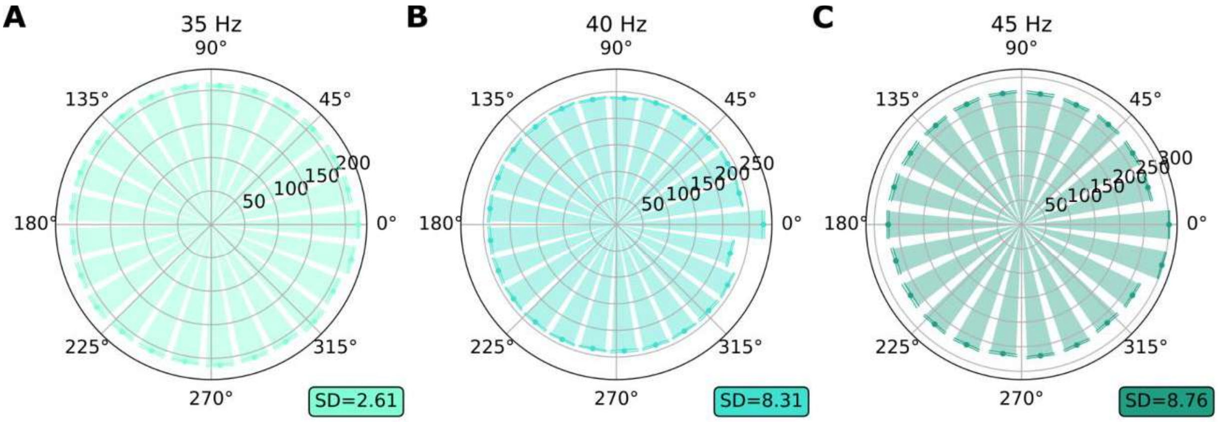


**Figure S2: TES-VS phase distributions are even across conditions.**
Phases between 39.9 Hz TES and VS in degrees. SD = standard deviation of the bins; bins in milliseconds. **A:** VS at 35 Hz. **B:** VS at 40 Hz. **C:** VS at 45 Hz.

### Analyses not conducted

The following statistical tests were preregistered but not run because they would only have been meaningful if the data supported our hypothesis that frequency-matched TES could enhance SSVEP amplitudes in at least one condition. The excluded tests, with SSVEP amplitudes as the dependent variable, were the following:

- Experiment 1, polarity effects: A permutation test between $VS_{40}TES_{O2\text{-}Cz+}$ and $VS_{40}TES_{Cz\text{-}O2+}$;
- Experiments 1 and 2, test-retest reliability of online effects: A correlation between $VS_{40}TES_{O2\text{-}Cz+}$ – $VS_{40}TES_{OFF}PRE$ in Experiment 1 and $VS_{40}TES_{O2\text{-}Cz+}$ – $VS_{40}TES_{OFF}PRE$ in Experiment 2;
- Calculating inverse Bayes factors for expected null effects in Experiments 1 and 2 (this was not possible because there was no effect to model the alternative prior).

### SSVEPs by block, Experiments 1 and 2

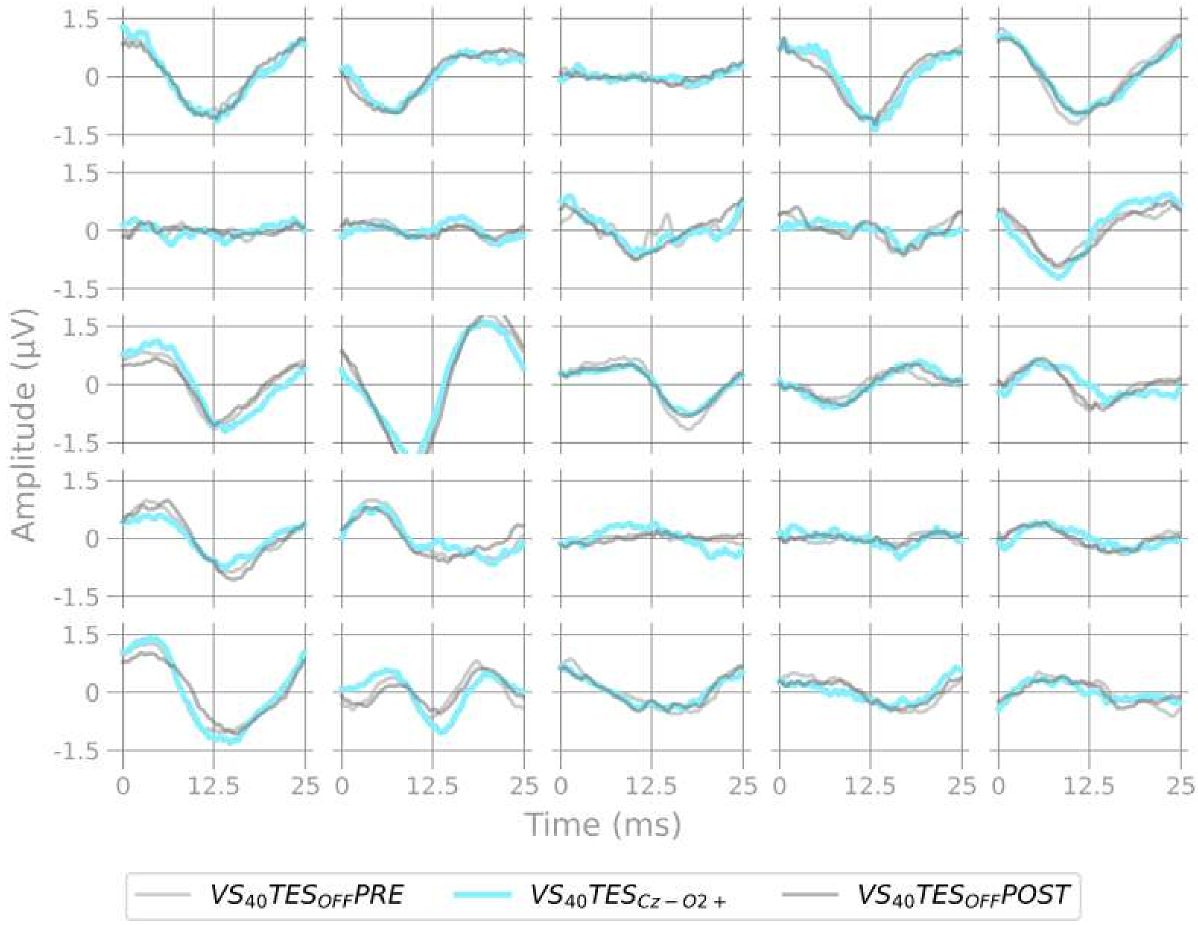


**Figure S3: SSVEPs Experiment 1, block Cz-O2+**

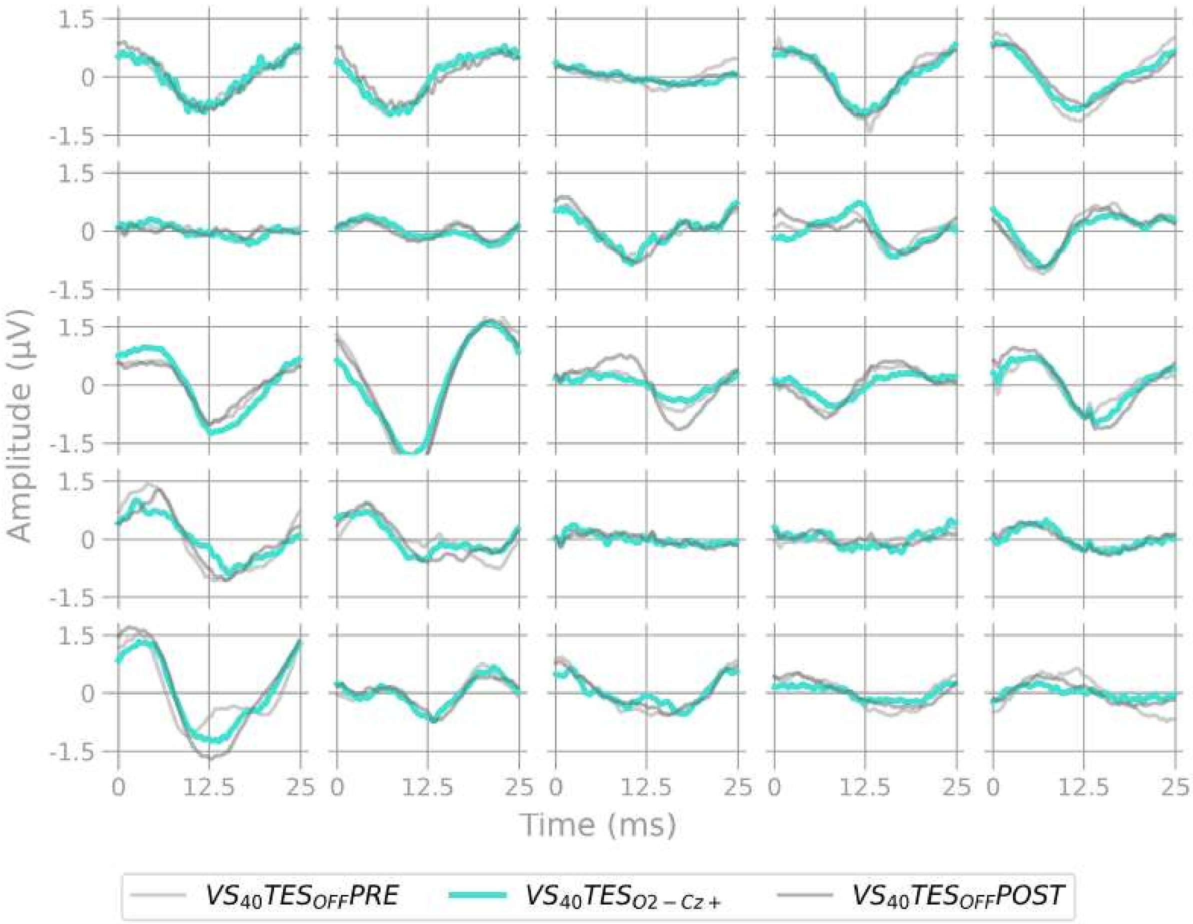


**Figure S4: SSVEPs Experiment 1, block O2-Cz+**

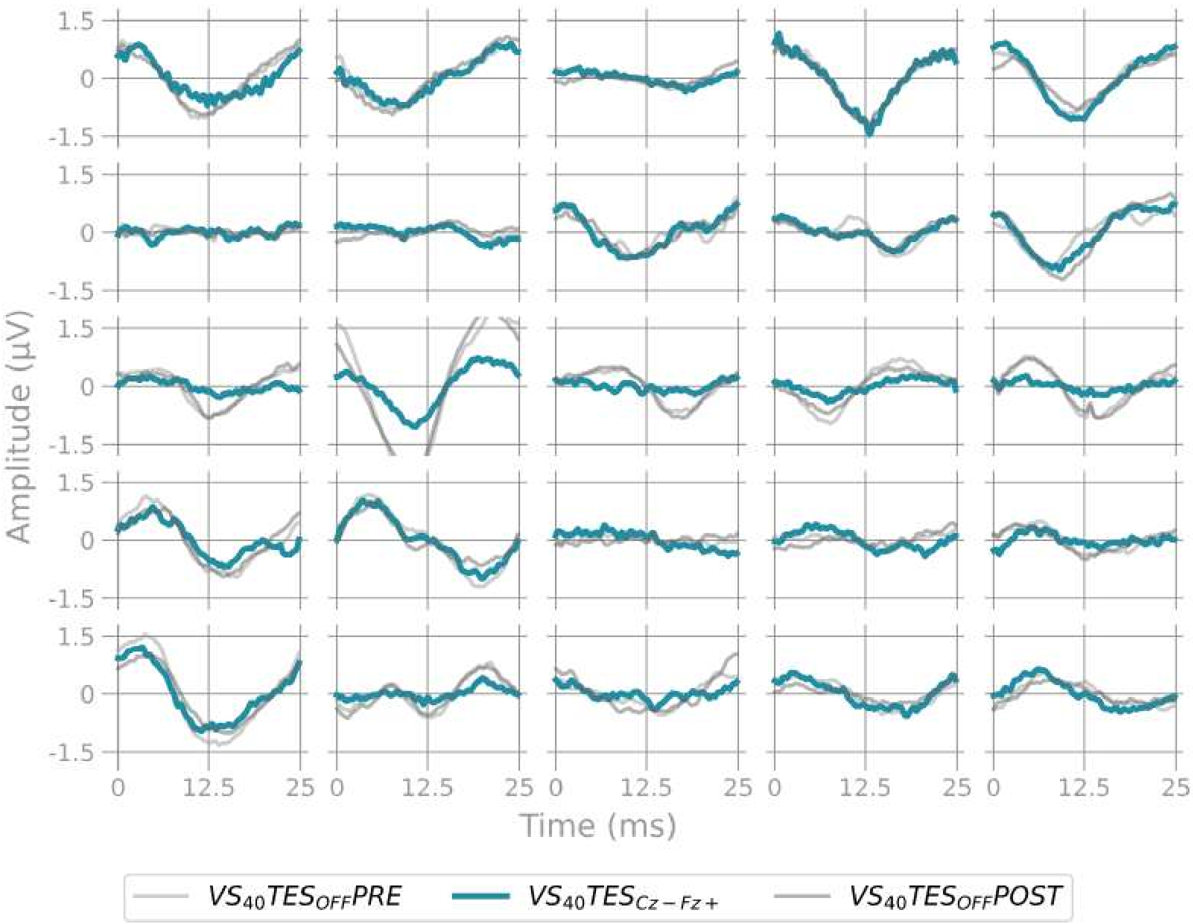


**Figure S5: SSVEPs Experiment 1, block Cz-Fz+**

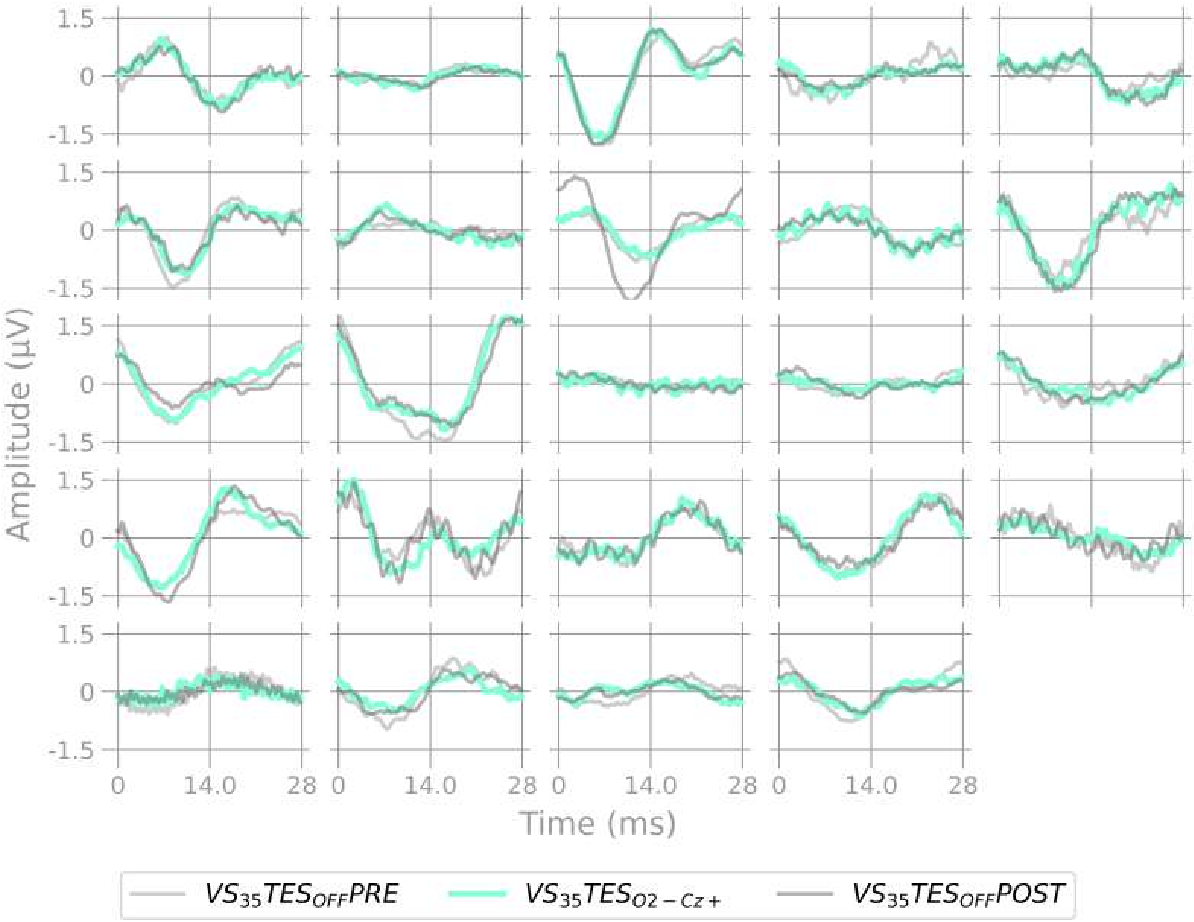


**Figure S6: SSVEPs Experiment 2, block 35 Hz**

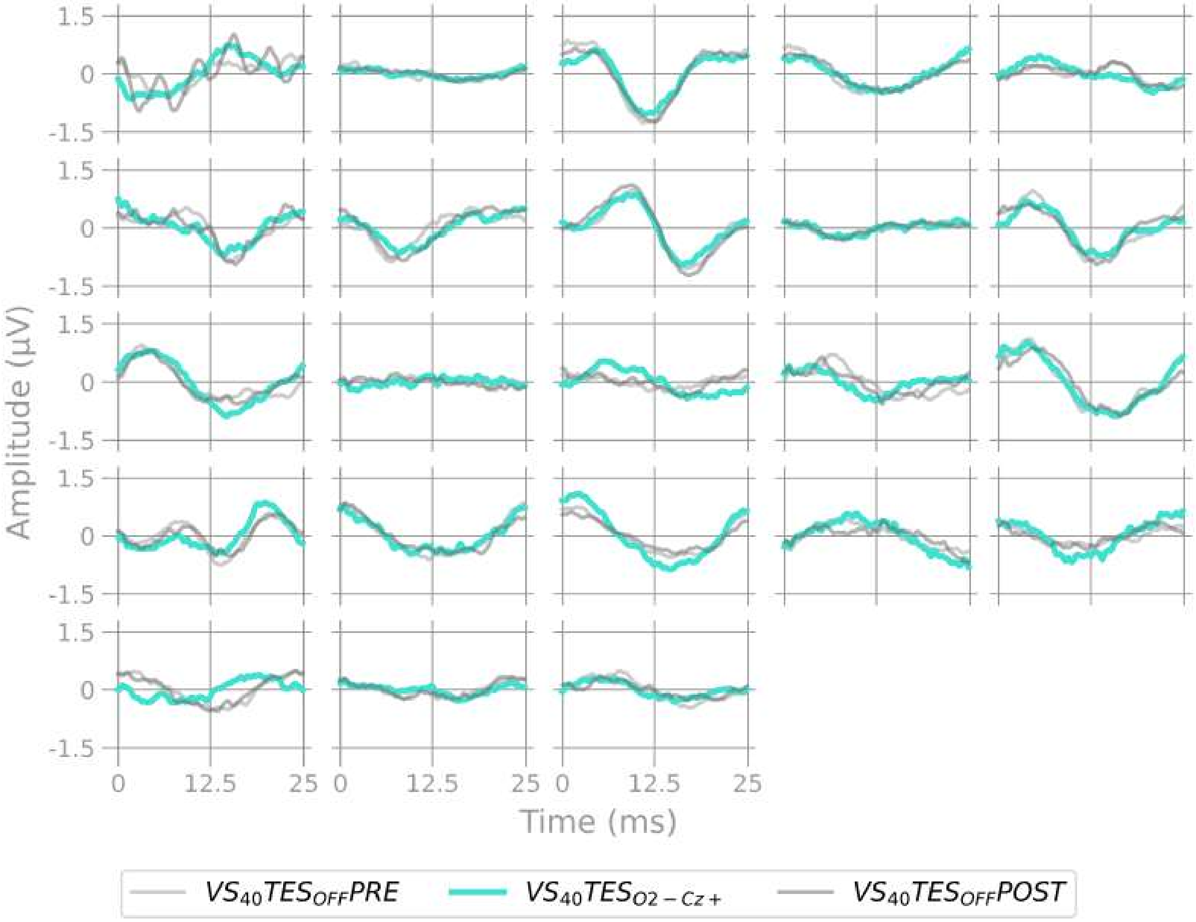


**Figure S7: SSVEPs Experiment 2, block 40 Hz**

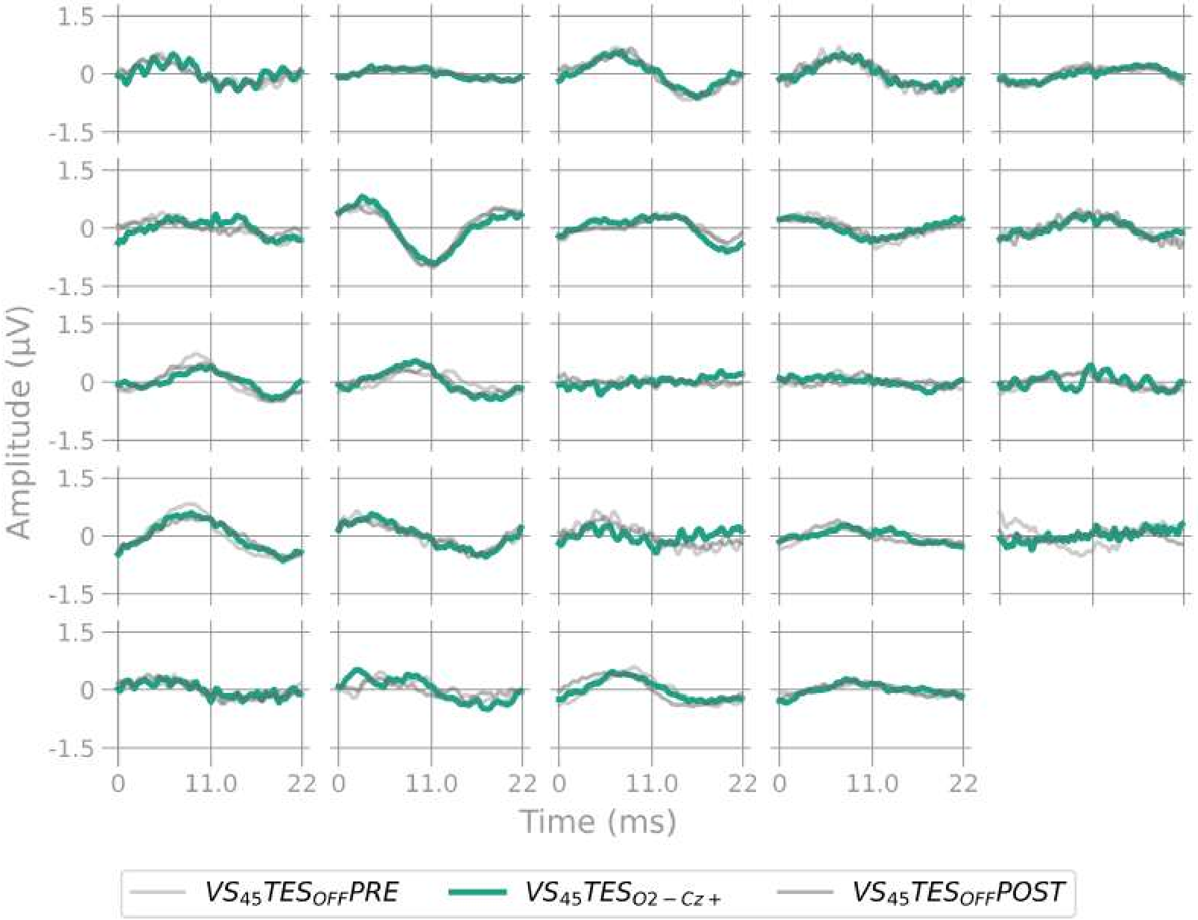


**Figure S8: SSVEPs Experiment 2, block 45 Hz**
